## Supplemental Fig 1 for "The endoplasmic reticulum-associated mRNA-binding proteins ERBP1 and ERBP2 interact in bloodstream-form *Trypanosoma brucei*"

```
P.parasit       MSTE------------EVVA-------VEEQEIPDVIERLPKPDKAEHEAKISALDGAIK  41
G.sulph         MANTT------------------SQGPTTQSQAMAVEQKLLTPE--ERQEEIRKLEQKKQ  40
ScBFR1          --------------------------------MSSQQHKFKRPDVSVRDKKLDTLNVQLK  28
Endotryp        -----------------------------------MPGALEKPKLSEYSAKISGLVEQRR  25
Paratryp        MSRSASAEAVATEPATTVVAEAPRRKPGQKPGWMSIPGGPKEPDRQAFSAKIREFGDRKR  60
Bodo            ----------------MAAVPAAENRPRTRPGWMNIPGGPKEPNFQDFRNKMQTLNTQKT  44
Blechomonas     ---------------MSAASPKEEHQRKRLPPWMSMPGAPPKPDFSAFGGKMAQLSAEKK  45
T.rangeli       MPGK---EIMSG---TADVAVATPAPSRRMPPWLLTPGGPKEPNTAEFRAKMAALAEEKR  54
T.cruzi         ----------MT---STEAAAVVAAPSKRMPPWLSMPGGPKEPNSAAFRAKMSALAEEKR  47
T.grayi         -----------------------------------MPGGPPEPNAAAFRSKMSRLAEEKR  25
T.theileri      --------MSAT---PPPAAAAAPTAKKQMPGWMSIPGGPPEPNGAAFRAKMNRLVEERR  49
T.vivax         -----------M---TSKVQTGAVEARRPPPRWMTGPDALPKPNVAEFRAKIAQLAKEKN  46
T.brucei        MSK-------TE---TAPEAAPAPTERRPQPRWMTAPGAPPRPDVMEHRTKMKALSQEKG  50
T.congolense    MGGD------AT---EVVAPAAGPAVRRGLPKWMTGPGAMPKPDFRAFKAKMAELAEEKR  51
                                                          *.      ::  :

P.parasit       KLQARTNVIRAEMDALKTNRGGYGGQIQ----------------------EAKAKF----  75
G.sulph         ELIQQIVVVQQPSKDELETAVAAQRAIIESALSQKEQKRLLQRQKYQKMEETKPEFLAAK  100
ScBFR1          KIDTEIGLIRKQIDQHQV-----------NDT----------------------------  49
Endotryp        ALLDELKQLQSTVQNDPE-----------RQK----------------------------  46
Paratryp        ALFDQLKEVQEKLHANGG-----------NEA----------------------------  81
Bodo            KLFDNLKELQGRAGPRDD---------TEREA----------------------------  67
Blechomonas     KLFEEIKRLQNSIRGDTS-----------NEA----------------------------  66
T.rangeli       TLLTEVRQLRASLGPREG-----------REA----------------------------  75
T.cruzi         ALLNEVKQLRASLGPREG-----------REE----------------------------  68
T.grayi         ALFTEVKQLRASLGPREG-----------REE----------------------------  46
T.theileri      ALLAEVKTLRASLGPREG-----------QEA----------------------------  70
T.vivax         TLFDKIKELRATLEPRDE------------------------------------------  64
T.brucei        ALIAQIKELRASLGPKSE------------------------------------------  68
T.congolense    SLIAQVKGLRASIGPKGD------------------------------------------  69
                 :  .   ::

P.parasit       -------AALRAEKDNLFQ-----------------QRNQITARLRQTRDEKDSTIKQQR  111
G.sulph         AAYEEQVTLLRKLREERKQLMGKIQETRKLMNDAQDQ---------NKSNNISTGNAALD  151
ScBFR1          ---------TQQERKKLQDKNKEIIKI---QADLKTRRSNIHDSIKQLDAQIKRKNNQIE  97
Endotryp        ---------CIAERNAFGEELNEIDACRKVQRELRAVQNAKIAKLRKSRTEVTEKLRSVQ  97
Paratryp        ---------IAVERQALRDRLGELEKERRGQRDVLTAKNEEIAALRKQRREIQDKQKELS  132
Bodo            ---------VMGERQELRRRMNEIDANRKKERDARSTKNEEISRIRRQRSDIEGKLKELS  118
Blechomonas     ---------LDEERKALRQRMGEIEAQRSAMRDLRVGKSEEISKFRKKRQETADKLRVLQ  117
T.rangeli       ---------TDNELAELRQRMSDIDARRKMEQEMRFKKNEEIQRVQKLHDERQSRLRELS  126
T.cruzi         ---------SGKELDGLRQRMGEIDNQRKVEQEMRFKKNEEIQRIQKLHDERVSRLRELS  119
T.grayi         ---------QENELGELRQRMGEIDTQRKAEQEMRFKKNEEIQKIQKVHEERLSKLRELS  97
T.theileri      ---------VEAELSELRQRMGEIDTQRKAEQEMRFKKNEEIQKIQKIHDERLNKLHELS  121
T.vivax         ---------NDKEREELRERIKELDNKKKLENDLRKKKSDEIQKVKKVQEEQLKKLRELT  115
T.brucei        ---------PDKERDEIRQRLKELDDKRKAEQEMRSKKSEEIIEVRKKHDEYVKKLRSLT  119
T.congolense    ---------NGKERDELFQRIKSIEEMRKAEQEMRSKKSSELNEAKKKHNENARKLRELT  120
                                                              .   :

P.parasit       -SVRANLKYGSVAEFDAAIAELKHKQETSSMSLNEEKRVIKEIEQLQAQKQQ-VSGFSDD  169
G.sulph         EK---LKNFKNLEQLDKHISSLERRQATESLSLTEEKKLVSEISLLKNKGRAFLENMQKL  208
ScBFR1          EKLGKKAKFSSTAEAKQRINEIEESIASGDLSLVQEKLLVKEMQSLNKLIKD-LVNIEPI  156
Endotryp        AE---VGGFTTLKEIDEAIDHMMRKMETSGGGLVAERRNQQHLQKLEEAKMH-LQKLQPL  153
Paratryp        TE---LGGFKTVQEIDEAIDYIMLKMETGGGGLSSEKKTAKRLHQLGEAKNL-LLQLQPL  188
Bodo            NE---LGAFRELSDIEMAIDHIMVRMETSGGGLASEKKAIKRLSQLEEAKSL-LLQLQPL  174
Blechomonas     AE---LGGFTDIAEIDTAIEFVMRKMETSSGGLAAEKRTIKRLQQLEEVKSL-LLQLQPL  173
T.rangeli       ED---LGGFTTLKEIDNAIALITRKMETSGGGLAAEKRTVRQLSKLEEAKRY-LVELQPL  182
T.cruzi         DD---LGGFTTIKEIDDAIAFVTRRMETSGGGLGAEKRAVRQLSKLEEAKRY-LVELQPL  175
T.grayi         DE---LGGFKSLKEIDGAIDYLTRKMETSSGGLAAEKRAVRQLNKLEEAKRY-LLELQPI  153
T.theileri      DG---LGGFKTLKEIDEAIAYLTRKMETSGGGLAAEKRTVRQLSKLEEAKRF-LLEMQPL  177
T.vivax         TE---LSGFKSVEEIDEAIAYMTKKMETSGGGLAAEKRMLRQLSQLEDAKRY-LQELQPV  171
T.brucei        DD---LGGFKSVEEFDEAIEYMTKKMETSGGGLAAEKRIMRQIGQLEDAKRY-LQELQPV  175
T.congolense    EE---LAGFKSVEEIDRAIVYMTRKMESSGGGLAAEKRMLQQLAQLEGAKRY-LQELQPL  176
                        :    : .  *  :     : . .*  *:    .:  *       :  :.
```

```
P.parasit    QGVVEKQNE-SIKEIRALQT---KKNEEIDAIQEKLNEQKQALDELYRL-----NEEENK  220
G.sulph      EAERKSSQEERKKELEKLFEEKKSLETKINETSKELERLKQSKDDIRTK-----QEISIA  263
ScBFR1       RKSV---DA-DKAKINQLKEELNGLNPKD--VSNQFEENQQKLNDIHSKTQGVYDKRQTL  210
Endotryp     TEAI---KE-ITEEEVILQQEYLAICEKIGILNGEYEEKLKQKNAKH-------KEAQED  202
Paratryp     DEAL---SE-FEEREFGLQQEYREIHERIGVINKEYEVQNDVRRK-----------ISDN  233
Bodo         QEAI---TD-ADDREASLQQEYREIHERIGALNKDYDEQYTVKQSKD-------KEAQKT  223
Blechomonas  TEAI---QE-AGDREVMLQQEHREIHERIGALNKKYEEELHTKQATE-------KNLHTT  222
T.rangeli    TEAI---TE-AKHREAMLQREWQEINDRIRSLWSEYNEQRATKQQKE-------QEIRIT  231
T.cruzi      TEAI---TE-AKHREAMLQREWQEINERIRNLRAEYNEQRATKQQKE-------QEIRSS  224
T.grayi      TEAI---TE-AKHREAMLQREWQEINERIRNLGTEYNEQRSMKQQRE-------QEMHST  202
T.theileri   TEAI---TE-AKHREALLQREWQEINERVRNLKSEYNEQRMTKQQKE-------QELRKT  226
T.vivax      NEAI---AE-AKHREALLQKEFDEINERIRGLTTACNEQRSTKMEKD-------QLVRGK  220
T.brucei     SEAI---AE-AKHCEATLQREIQEINERIRGLNEEYKGQRSTKMEKD-------DKMRST  224
T.congolense SEAV---DE-AKHHEATLLRELQEINERIRDLNSKYKDQRSVKMQKD-------EEIRSA  225
                              *         .        .

P.parasit    KDKFPALAKERKEIKEQLDEKFTAIKTLRKEFKEANDKYYNNIRLVRKKKELERQKEEEA  280
G.sulph      KISEELSEVNLDDIQKRIDAANEEIRRLRQDFNQKLNKWYNYKKRSSE-------LDRLN  316
ScBFR1       FNKRAALYKKRDELYSQIRQI-------RADFDNEFKSFRAKLDKER------LKREEEQ  257
Endotryp     GAHRAEVYKKCDALRVRIAEISQTVESLRAERDRLSSEWHAWSREARTKYYAQLEQRREQ  262
Paratryp     IVDKKPLFERRDLIRKELDAVSVEMDTLRAAHRQAEEAWDKWRVEAQAKYAAKIKAEREE  293
Bodo         SVDRTVLIKERDDLRQKITKLNEEMTKLREGFNTEKESWEAWREEAIKKYGEKMEAERKE  283
Blechomonas  FQQRTSVYKQCDELREKINKLNEEMDTYRASHNDAVQKWETWCAEAREKYTAKMETERQE  282
T.rangeli    GVNRSEVYKKCEEISAKINKLSEEMNRMREEHNKAMEAWNAWREEARAKYLAKMEAERKE  291
T.cruzi      GVNRAEIYKKCEEINGKITKLSGEMDRLREEHNKAMEAWNTWREEARAKYIAKMEEERKE  284
T.grayi      GANRAEVYKKCDAISAKINKLNEEMNTLREEHNKALEAWNAWREEARTKYAAKVEAERKE  262
T.theileri   GVNRQEVYKKCDEISAKIKSLSDEMNTLREEHNKAVEAWNAWREEARAKYAAKVEAERKE  286
T.vivax      GASRQEVYAKCSELHAKITGIVEKMNTLREEHSKAMEAWNAWRAEAFAKHQAKMEEIRKE  280
T.brucei     GANRQEVFKKCDEISAKITSITKEMNALSEDFQKAMEVWKSWCDEARAKHMAKMEEVRKE  284
T.congolense GANRQEVFKKCDEINANIAKIAQEMNSLSEAHQKAMEKWNAWCDEARAKHIAKMEEIQKD  285
                      . . :  .:                  . :                  .

P.parasit    RKAEY--E------AKL--ADYEKEMAKIHPYQDEMDLCDALVSFLEKTYAKELKEEQDE  330
G.sulph      RHLRFQYRQYQRDLKRVEKEKELAEYGTPDPYEEEKAMCDNLIQ-----YLQRLSKIDVD  371
ScBFR1       RLSKLLEQKDV-DMGKL-QEKLTHAKI--PAFTYEIGAIENSLLVLDPTYVKPKKNILPD  313
Endotryp     RRKEYEEIRN---AHKI-AAKRERAAKRQNPYVTEISACATLIQ-----YLKQKKVMLEQ  313
Paratryp     KLRRYLERRD---ASKL-AEKRERAMKRMNPYASEIASCITLVQ-----YLRDKRMMSQR  344
Bodo         RERRRNEYMN---AEKI-ARKQARATKRQNPHETQIGACSTLVR-----YLRDRIVMSQR  334
Blechomonas  RYKRELERHT---NAKL-TEKLNRAQRRRNPYEMEISACDTLSQ-----YLIDKKHMVTR  333
T.rangeli    RQRRYLERKN---AAKL-EEKRARALRRQNPYEVEIEACKTLLR-----YVQDHKVMVQR  342
T.cruzi      RRRRYLERKN---AAKI-EAKRARALRRQNPYEAEIDACKTLLR-----YMQDQKVMVQR  335
T.grayi      RHLRYLERKN---AAKL-EEKRARAMRRQNPYEAEVDACSVLVR-----YLRDHKMMVQR  313
T.theileri   RQRRYLEYKN---AAKM-EEKRARALRRQNPYEAEADACSTLVR-----YLRDKKAMVQR  337
T.vivax      RERRIHEKNN---AAKL-EEKRARALRRMNPYEVEIAACDTLIQ-----YLREQKVMVKR  331
T.brucei     RHRRFLERKN---APKL-AEKRERALRRMNPYEVEIAACDTLLQ-----YLRDQKIMVQR  335
T.congolense RQRRIMERKN---AAKL-EEKRARALRRMNPYEVELAACDTLLR-----YLGEQKIMVQR  336
             :  .           ::   .          .  :              *

P.parasit    K---------AAETTAAPLE-LDGMKPLQRKEEDFMMLGGGKKGK--KGRNGK-------  371
G.sulph      ENKKTKKDVQNIE--------LNGAKLIGKNASFFEAEPNTNVSKG--------------  409
ScBFR1       LS-----SN-ALETKPARKVVADDLVLVTPKKDDFVNVAPSKSKKYKK-KNQQKN-----  361
Endotryp     EEQERKKREAAAHFDPSQTA-PAGCVVLNDSKWADNKTPYKSTTKLPKQQKKQKEKIPQA  372
Paratryp     EEEERVKCEAAAKFDPARAA-PSGFVMLGEDKWSSPHSAPKKGGK----KQSAKA--PSA  397
Bodo         DEEERKRRVAMASFDPSASA-PSGFALAAPIELPKKKSKAA-------------------  374
Blechomonas  EEEERVKREAAATFDPAKAV-PAGCVVLNEGKWGK-NAKPAPKAS----KKQQKT-NTAP  386
T.rangeli    EEAELARKQAAAAFDPSKFL-PEGAVLLNDGKKFSDSRKGGAGGK----HKAAQT----Q  393
T.cruzi      EEVELARKRAAATFDPTNFL-PEGAVLLNDGKKFSEPHKAAPGGK----SKTKQN----Q  386
T.grayi      EEEELARKNAAATFDPTKFL-PEGAVVLNDGKKWTEPHKAAVGGK----KNKQQQKQQQQ  368
T.theileri   EEEELARQEAAATFDPTKFL-PEGAVLLNDGKKRTDANKGGALGK----KNKQQQ----Q  388
T.vivax      EEEERIKKEAVANFDPAKFA-PSGAVILNDGKNWKDHAKGAVGGK----NKQR------P  380
T.brucei     ENEERARREAAANFDPAEFA-PEGAVVLNDGMSHQNG---GDSKK----QKQQ------A  381
T.congolense ENEERAKREAAAAFDPTKFA-PEGAVLLNDGKGAGEG---KSQKS----QH-T------K  381
                                    .
```

```
P.parasit      ------KTKKASKLVLPL----------AQMEAFSTIGLLPPASAAAVSESLAAVKTKKV  415
G.sulph        RKTKRKGAAKDSTFSKAVDSEKLPPHNMEYFLAFQKLNVVPPVYVKDILGTIELLKERKS  469
ScBFR1         ----TENEQPASIFNKVDGKFTLEP---TLIATLAELDVTVPINSDDVKITVEQLKKKHE  414
Endotryp       KAGTTLSDQKERPLHHTDEK----------IRLFRIIEIEPTRSRVAIDSTISEIESAKK  422
Paratryp       KTTEAPVATKDRVLQHSEEK----------QQMFQAVRVDPPLALSAIDAAIKAIEERRK  447
Bodo           ---PKAEDKTERTVTHNDEK----------KRLFASVGISAPATLSEVEKTIEQLKKKQA  421
Blechomonas    PVTAAKRVDSTRMLQHPEEK----------MKLFHLIDLEPPITITAFDSTIQAIKAKRK  436
T.rangeli      KQKPEK-APKNRVLQHPEDK----------IRLFQLVNEELPVALAAIDETMERLRAKQK  442
T.cruzi        KQKSESAPPKNRVLQHPEDK----------IRLFQLINEEPPVALSAIDGAMETIRSKQK  436
T.grayi        QQKQQKTEPKNRVLQHGEEK----------IRLFQLIGEEPPLALAAIDDAVKRISAKQK  418
T.theileri     QQKPKAGTAKNRVLQHSEDK----------IRLFQLINEEPPLALAAIDASVERIRAKQV  438
T.vivax        TSKKESGASKAVSIKHGEEK----------VELFTLIGEEPPKVLDDIDTLLTKITEKRK  430
T.brucei       NKAKRDSAPKPRVIKHSEEK----------LELFKLVDEQPPRFLEDIGGIMENIRAKLK  431
T.congolense   KVNGTESAAKSGVIKHSDEK----------VKLFKLVSEEPPQSVGDIDGVMESLRTKQQ  431
                            .                   :  :          .   :  :

P.parasit      WFNEQTSRPK------------AGKVVEPAAEAAP--------AKVVSPKKKSSKNNKFN  455
G.sulph        YYENAPE---------------KVDLEATEEEDLS-------------TLLPKSDSDNVI  501
ScBFR1         ELLSKQEEQTKQNIESVEKEIEKLNLDYSNKEQQV--KKELEE-KRLKE-------QEES  464
Endotryp       KYESHIQ---------------TGELVLSSGEDEEDDEENTVNDDEVPSELAPADAQGVV  467
Paratryp       VMETHIV---------------TGEPVLSSDDENEEENAEETNSPVA---------NGDE  483
Bodo           EYESHIK---------------TGDLVLSSDDEEEEKEEAAPADE---------------  451
Blechomonas    EYESHIT---------------TGDIVLSSESDGEGETHPEDSNLAEPDAAPA-------  474
T.rangeli      EYESHIK---------------VGELELSSDDEEDEEAQEEEG--AAEATEE----E-AP  480
T.cruzi        EYESHIK---------------TGELELSSDDEEEEEEQPQEEEETAAAATE----E-VQ  476
T.grayi        EYESHIK---------------TGDLELSSDDEEEEEQEEPQPQEEKGEEQEV--VDGAK  461
T.theileri     EYESHKK---------------TGELELSSDDDEEEEEEENVEQEQKEEQNQGDEEQGED  483
T.vivax        EYASHIK---------------VGELELSSDDDEQEHPQEEEEEAEVEDAVESSKEKGEK  475
T.brucei       EYSSHIK---------------TGEPELSSDDEEDEEKKEQEGEEEAVA-------KEDE  469
T.congolense   EYASHIK---------------TGEPELSSDDEEEQEQEQEQEQEQQEN-------EGAA  469
                                        .      .

P.parasit      ASDKDAFPSLGGVAAELPSWGPGMAPAVAEPAVAAEFAEAD---VVTESE-   502
G.sulph        VEDREDFPDG------LPIVSNGYS-YRAQQGSKPSFAEVMQHSSLTDSVD   545
ScBFR1         EKDK------------------EN---------------------------   470
Endotryp       VDDA----------------KKDIAE-------------------------   477
Paratryp       KEEA---EEA------APVLAAA----------------------------   497
Bodo           ---------------------------------------------------   451
Blechomonas    EHNE----EP------VPEVTAE----------------------------   487
T.rangeli      KEAD----DV------EE-VTAGAAEAVTQSGQEEATV-------------   507
T.cruzi        KKKD----EV------EKEVTADKIGLVTQ---NEETVEA-----------   503
T.grayi        TESQ----LE------ESEVKADE---------------------------   475
T.theileri     KE-------------------------------------------------   485
T.vivax        EEEK----EG------EEKKDEERAE-------------------------   491
T.brucei       TNEG----EG------EDFA-------------------------------   479
T.congolense   EEEG----DT------N----------------------------------   476
```

| **Abbreviation** | **Species** | **Gene ID** |
|---|---|---|
| P.parasit | *Phytophthora parasitica* | ETI45734 |
| G.sulph | *Galderia sulphararia* | XP_005709255 |
| ScBFR1 | *Saccharomyces cerevisiae* | BFR1 |
| Endotryp | *Endotrypanum moterogeii* | EMOLV88_320005100 |
| Paratryp | *Paratrypanosoma confusum* | PCON_0018750 |
| Bodo | *Bodo saltans* | BSAL_12610 |
| Blechomonas | *Blechomonas ayalai* | rna_Baya_154_0090 |
| T.rangeli | *Trypanosoma rangeli* | TRSC58_05341 |
| T.cruzi | *Trypanosoma cruzi* | TcCLB.506525.10 |
| T.grayi | *Trypanosoma grayi* | DQ04_02291030 |
| T.theileri | *Trypanosoma theileri* | TM35_000033410 |
| T.vivax | *Trypanosoma vivax* | TvY486_1013600 |
| T.brucei | *Trypanosoma brucei* | Tb927.10.14150 |
| T.congolense | *Trypanosoma congolense* | TcIL3000_10_12030 |

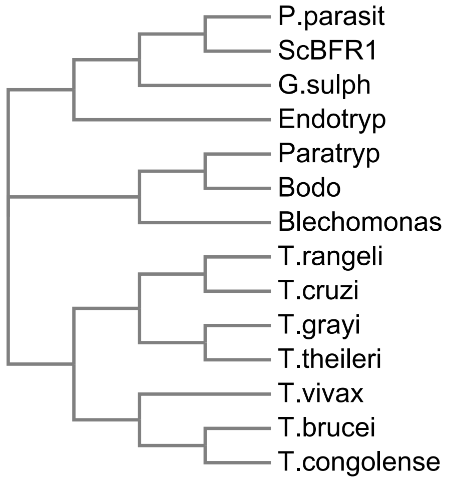
