## Supplementary figures and images for "The endoplasmic reticulum-associated mRNA-binding proteins ERBP1 and ERBP2 interact in bloodstream-form *Trypanosoma brucei*"

### Supplemental Fig 2

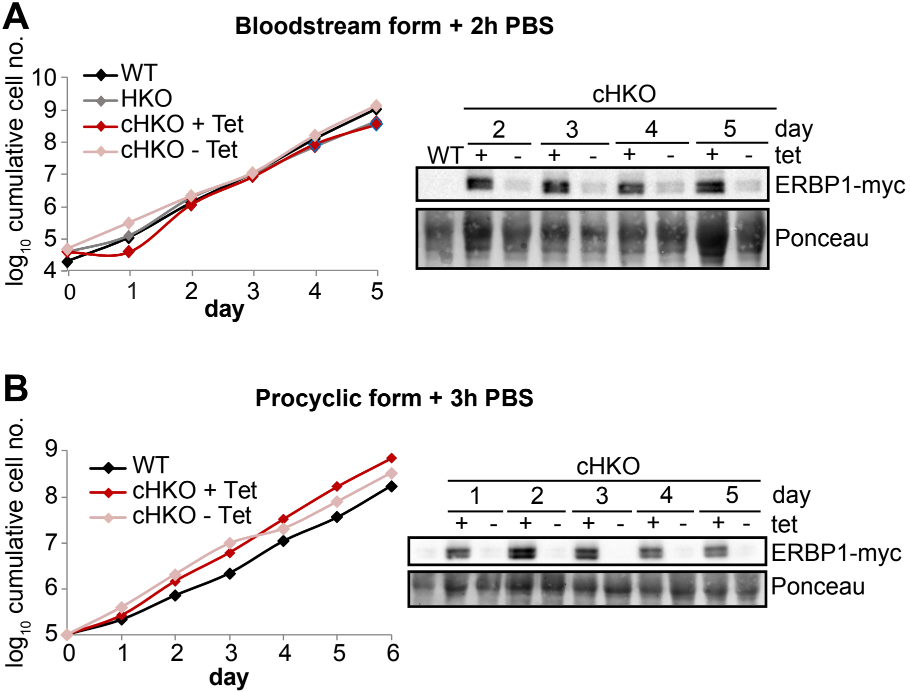

### Supplemental Fig 3

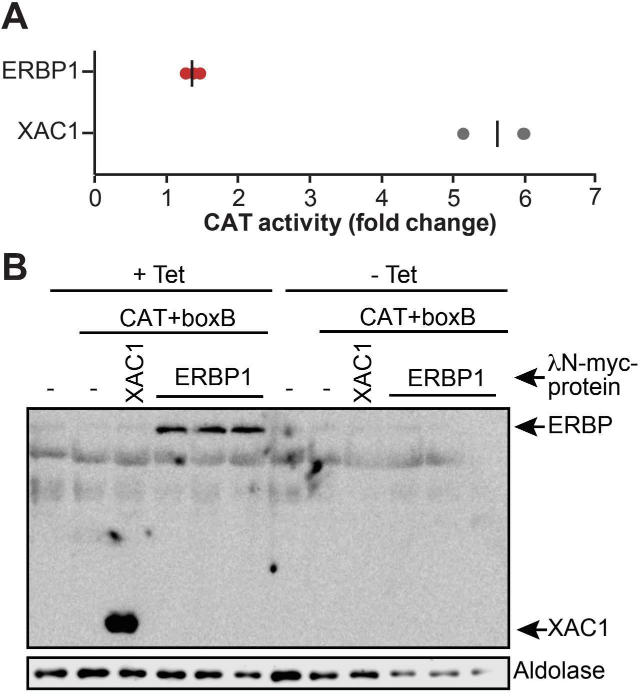
